# Simulations of biomass dynamics in community food webs

**DOI:** 10.1101/070946

**Authors:** Eva Delmas, Ulrich Brose, Dominique Gravel, Daniel B. Stouffer, Timothée Poisot

## Abstract

1. Food webs are the backbone upon which biomass flows through ecosystems. Dynamical models of biomass can reveal how the structure of food webs is involved in many key ecosystem properties, such as persistence, stability, etc.
2. In this contribution, we present `BioEnergeticFoodWebs`, an implementation of Yodzis & Innes (1992) bio-energetic model, in the high-performance computing language Julia.
3. We illustrate how this package can be used to conduct numerical experiments in a reproducible and standard way.
4. A reference implementation of this widely used model will ease reproducibility and comparison of results across studies.

## INTRODUCTION

Community and ecosystem ecologists have long sought to understand the diversity, properties, and dynamics of multi-species assemblages. The characteristics of communities emerge in unpredictable ways because species influence one another through direct, and indirect, ecological interactions. Seeing that the coexistence of populations is constrained at least by feeding interactions, models of the relationship between resources and consumers have provided a useful and frequent tool in studying the theory of community dynamics. Although these modeling efforts started from simple, abstract models like those from the Lotka-Volterra family (Bacaër 2011), more tailored and parameterized models have emerged whose goal was to include a broader range of ecological and biological mechanisms, thus hopefully providing more realistic representations of empirical systems. Among these, the “bio-energetic” model of Yodzis & Innes (1992) is a general representation of resource-consumer dynamics, yielding results comparable to empirical systems, while needing minimal parameters. To achieve this purpose, it uses allometric scaling of metabolic biomass production and feeding rates, meaning that the flow of biomass from a resource to its consumer depends on their body mass.

Since the work of Yodzis & Innes (1992), Chesson & Kuang (2008) have shown that the dynamics of ecological communities are driven not only by pairwise interactions, but also by the fact that these interactions are embedded in larger networks, and Berlow et al. (2004) show how disturbances affecting species biomass or density cascade up, not only to the species that they interact with, but with species up to two degrees of separation from the original perturbation. In this context, models of energy transfer through trophic interactions are better justified when they account for the entire food-web structure, such as Williams et al. (2006) adaptation of Yodzis & Innes (1992) model. This food-web bio-energetic model has been used, for example, to show how food web stability can emerge from allometric scaling (Brose et al. 2006b) or allometry-constrained degree distributions (Otto et al. 2007) (more past uses of the model are described in supplementary table S1). Yet, although these and other studies used the same mathematical model, implementations differ from study to study and none have been released. Motivated by the fact that this model addresses mechanisms that are fundamental to our understanding of energy flow throughout food webs, we present `BioEnergeticFoodWebs` (Bio-Energetic Food-Webs Model), a *Julia* package implementing Yodzis & Innes (1992) bio-energetic model adapted for food webs (Williams et al. 2006) with updated allometric coefficients (Brown et al. 2004; Brose et al. 2006b).

This package aims to offer an efficient common ground for modeling food-web dynamics, to make investigations of this model easier, and to facilitate reproducibility and transparency of modeling efforts. Taking a broader perspective, we argue that providing the community with reference implementations of common models is an important task. First, implementing complex models can be a difficult task, in which programming mistakes will bias the output of the simulations, and therefore the ecological interpretations we draw from them. Second, reference implementations facilitate the comparison of studies. Currently, comparing studies means not only comparing results, but also comparing implementations – because not all code is public, a difference in results cannot be properly explained as an error in either studies, and this eventually generates more uncertainty than it does answers. Finally, having a reference implementation eases reproducibility substancially. Specifically, it becomes enough to specify which version of the package was used, and to publish the script used to run the simulations (as we do in this manuscript). We fervently believe that more effort should be invested in providing the community with reference implementations of the models that represents cornerstones of our ecological understanding.

## THE MODEL

### Biomass dynamics

We implement the model as described by Brose et al. (2006b), which is itself explained in greater detail in Williams et al. (2006). This model describes the flows of biomass across trophic levels, primarily defined by body size. It distinguishes populations based on two variables known to drive many biological rates: body mass (*i.e*. how large an organism is, how much biomass it stocks) and metabolic type (*i.e*. where the organism get its biomass from and how it is metabolized). Once this distinction made, it models populations as simple stocks of biomass growing and shrinking through consumer-resources interactions. The governing equations below describe the changes in relative density of producers and consumers respectively.

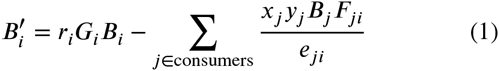

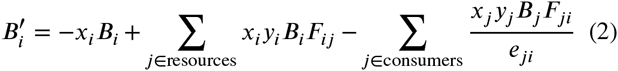

where *B*_*i*_ is the biomass of population *i, r*_*i*_ is the mass-specific maximum growth rate, *G*_*i*_ is the net growth rate, *x*_*i*_ is *i*’s mass-specific metabolic rate, *y*_*i*_ is *i*’s maximum consumption rate relative to its metabolic rate, *e*_*ij*_ is *i*’s assimilation efficiency when consuming population *j* and *F*_*ij*_ is the multi-resources functional response of *i* consuming *j*:

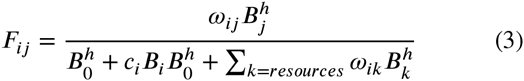

### Growth rate function

The formulation of the growth rate *G*_*i*_ can be chosen among three possibilities (Williams 2008) that all share the general equation of *G*_*i*_ *=* 1 – *s/k*, where *s* is the sum of biomass of populations in competition for a ressource with carrying capacity *k*. The first scenario, used by Brose et al. (2006b), sets *s = B*_*i*_ and *k = K*: species only compete with themselves for independent resources. The issue with this formulation (Kondoh 2003) is that the biomass and productivity of the system scales linearly with the number of primary producers. The second formulation “shares” the resource across primary producers, with *s = B*_*i*_ and *k = K/n*_*P*_, wherein *n*_*p*_ is the number of primary producers. Finally, a more general solution that encompasses both of the previous functions is *s* = ∑*α*_*ij*_*B*_*j*_, with *α*_*ii*_ (intraspecific competition) set to unity and *α*_*ij*_ (interspecific competition) taking values greater than or equal to 0. Note that *α*_*ij*_ = 0 is equivalent to the first scenario of *k = K* and *s* = *B*_*i*_.

### Numerical response

In Equation 3, *ω*_*ij*_ is *i*’s relative consumption rate when consuming *j*, or the relative preference of consumer *i* for *j* (McCann et al. 1998; Chesson & Kuang 2008). We have chosen to implement its simplest formulation: *ω*_*ij*_ *= l/n*_*i*_, where *n*_*i*_ is the number of resources of consumer *j*. The Hill coefficient *h* is responsible for the hyperbolic or sigmoidal shape of the functional response (Real 1977), *B*_0_ is the half saturation density and *c* quantifies the strength of the intraspecific predator interference – the degree to which increasing the predator population’s biomass negatively affect its feeding rates (Beddington 1975; DeAngelis et al. 1975). Depending on the parameters *h* and *c* the functional response can take several forms such as type II (*h* = 1 and *c =* 0), type III (*h* > 1 and *c =* 0), or predator interference (*h* =1 and *c >* 0).

### Metabolic types and scaling

As almost all organisms’ metabolic characteristics vary predictably with body mass (Brown et al. 2004), these variations can be described by allometric relationships as described in Brose et al. (2006b). Hence, the per unit biomass biological rates of production, metabolism and maximum consumption follow negative power-law relationships with the typical adult body mass (Savage et al. 2004; Price et al. 2012).

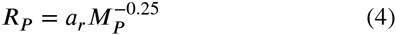

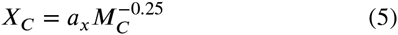

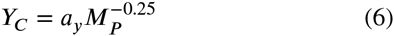

Where the subscripts P and C refer to producers and consumers populations respectively, M is the typical adult body mass, and *a*_*r*_*, a*_*x*_ and *a*_*y*_ are the allometric constant. To resolve the dynamics of the system, it is necessary to define a timescale. To do so, these biological rates are normalized by the growth rate of the producers population (*cf*. Equation 4) (Brose et al. 2006b; Williams et al. 2006).

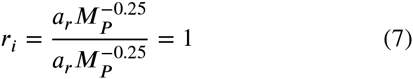

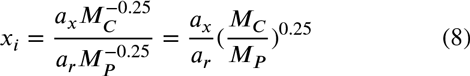

In Equation 1 and Equation 2, *y*_*i*_ refer to the maximum consumption rate of population *i* relative to its metabolic rate and thus become a non-dimensional rate:

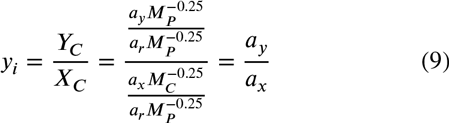

As the biological rates also vary with the organisms metabolic types, the maximum consumption rate of population *i* relative to its metabolic rate (*y*_*i*_) is not the same for ectotherm vertebrate (*y*_*i*_ *=* 4) and invertebrate (*y*_*i*_ = 8) predators; the same goes for the allometric constant *a*_*x*_, which causes the mass-specific metabolic rate (*x*_*i*_) to differ for ectotherm vertebrates (*a*_*x*_ *=* 0.88) and invertebrates (*a*_*x*_ *=* 0.314). The diet of predators also affects their assimilation efficiency (*e*_*ij*_) which is greater for carnivores (*e*_*ij*_ *=* 0.85) than for herbivores (*e*_*ij*_ *=* 0.45).

Based on the observation that most natural food webs have a constant size structure (Brose et al. 2006a; Hatton et al. 2015), the consumer-resource body-mass ratio (*Z*) is assumed to be constant. The body mass of consumers is then a function of their mean trophic level (*T*), it increases with trophic level when *Z* ≥ 1 and decreases when *Z ≤* 1:

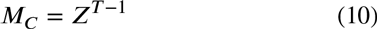

where *M*_*c*_ is the body mass of consumers, normalized by the body mass of the basal species (*T =* 1) to make the results independent of the body mass of the basal species.

### Setting the simulation parameters

All of these parameters can be modified before running the simulations (see `?model_parameters`), and are saved alongside the simulation output for future analyses. The default values and meanings of the different parameters are explained in the documentation of the `model_parameters` function. The user can specify which species are ectotherm vertebrates by supplying an array of boolean values, and the body mass of each species by supplying an array of floating-point values.

### Saving simulations and output format

The core function `simulate` performs the main simulation loop. It takes two arguments, `p` – the dictionary generated through the `model_parameters` function and containing the entire set of parameters – and `biomass`, a vector that contains the initial biomasses for every population. Three keywords arguments can be used to define the initial (`start`) and final (`stop`) times as well as the integration method (`use`, see `?simulate` or the online documentation for more details on the numerical integration methods available). This function returns an object with a fixed format, made of three fields: `:p` has all the parameters used to start the simulation (including the food web itself), `:t` has a list of all timesteps (including intermediate integration points), and `:B` is is a matrix of biomasses for each population (columns) over time (rows). All measures on output described below operate on this object.

The output of simulations can be saved to disk in either the `JSON` (javascript object notation) format, or in the native `jld` format. The `jld` option should be preferred since it preserves the structure of all objects (`JSON` should be used when the results will be analyzed outside of `Julia`, for example in `R`). The function to save results is called `BioEnergeticFoodWebs.save` (note that the prefix `BioEnergeticFoodWebs`. is mandatory, to avoid clashes with other functions called `save` in base `Julia` or other packages).

### Measures on output

The `BioEnergeticFoodWebs` package implements a variety of measures that can be applied on the objects returned by simulations. All measures take an optional keyword argument `last`, indicating over how many timesteps before the end of the simulations the results should be averaged.

Total biomass (`total_biomass`) is the sum of the biomasses across all populations. It is measured based on the populations biomasses (`population_biomass`).

The number of remaining species (`species_richness`) is measured as the number of species whose biomass is larger than an arbitrary threshold. Since `BioEnergeticFoodWebs` uses robust adaptive numerical integrators (such as ODE45 and ODE78) the threshold default value is *ε, i.e*. the upper bound of the relative error due to rounding in floating point arithmetic. In short, species are considered extinct when their biomass is smaller than the rounding error. For floating point values encoded over 64 bits (IEEE 754), this is around 10^−16^. An additional output related to `species_richness` is `species_persistence`, which is the number of persisting species divided by the starting number of species. A value of `species_persistence` of 1 means that all species persisted. A value of `species_persistence` of 0 indicates that all species went extinct.

Shannon’s entropy (`foodweb_evenness`) is used to measure diversity within the food web. This measure is corrected for the total number of populations. This returns values in]0; 1], where 1 indicates that all populations have the same biomass. It is measured as

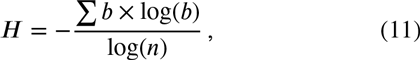

where *n* is the number of populations, and *b* are the relative biomasses (*b*_*i*_ *= B*_*i*_/∑ B).

Finally, we used the negative size-corrected coefficient of variation to assess the temporal stability of biomass stocks across populations (Tilman 1995) (`population_stability`). This function accepts an additional `threshold` argument, specifying the biomass below which populations are excluded from the analysis. For the same reason as for the `species_richness` threshold, we suggest that this value be set to either the machine’s *ε*(0.0) (*i.e*. the smallest value immediately above 0.0 that the machine can represent), or to 0.0. We found that using either of these values had no qualitative bearing on the results described below. Values close to 0 indicate little variation over time, and increasingly negative values indicate larger fluctuations (relative to the mean standing biomass).

## IMPLEMENTATION AND AVAILABILITY

The `BioEnergeticFoodWebs` package is available for the `julia` programming language, and it is continuously tested on the current version of `Julia`, the release immediately before and on the current development version. `Julia` is an ideal platform for this type of models, since it is easy to write, designed for numerical computations, extremely fast, easily parallelized, and has good numerical integration libraries. The package can be installed from the `Julia` REPL using `Pkg.add(“BioEnergeticFoodWebs”)`. A user manual and function reference is available online at http://poisotlab.io/BioEnergeticFoodWebs.jl/latest/, which also gives instructions about installing Julia, the package, and how to get started.

The code is released under the MIT license. This software note describes version 0.2.0. The source code of the package can be viewed, downloaded, and worked on at https://github.com/PoisotLab/BioEnergeticFoodWebs.jl. Potential issues with the code or package can be reported through the *Issues* system or at https://gitter.im/PoisotLab/BioEnergeticFoodWebs.jl. The code is version-controlled, undergoes continuous integration, and has a code coverage of approx. 90% to this date.

**Figure 1.**
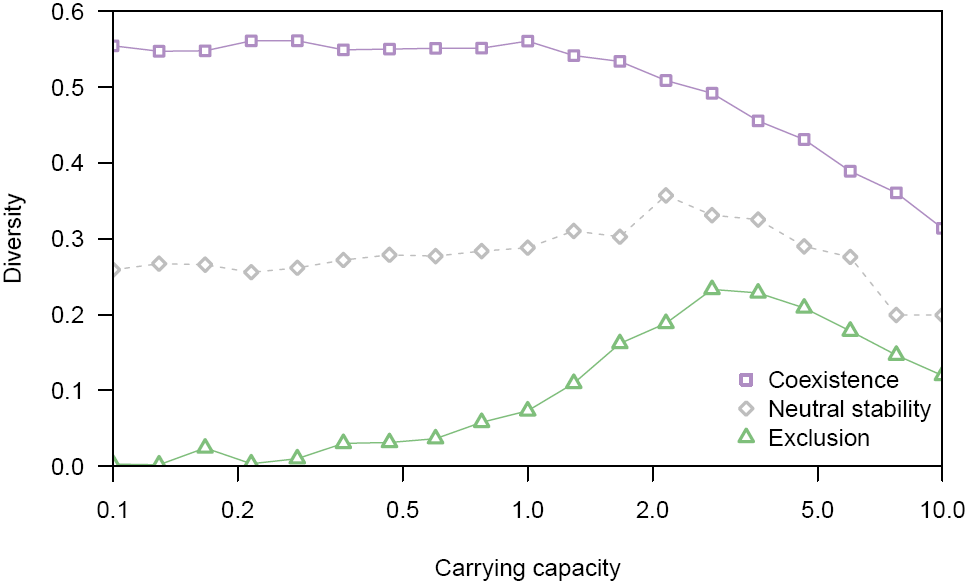
Effect of increasing the carrying capacity of the resource for different levels of competition (*α* ∈ [0.9,1.1]). For conditions of neutral coexistence or coexistence (*α ≤* 1), diversity is stable until *K* ≈ 5. For conditions of competition exclusion (*α* > 1), diversity increases for *K <* 5, and decreases after.

## USE CASES

All functions in the package have an in-line documentation available at http://poisotlab.io/BioEnergeticFoodWebs.jl/latest/, as well as from the `julia` interface by typing ? followed by the name of the function. In this section, we will describe three of the aforementionned use cases. The code to execute them is attached as Supp. Mat. to this paper. As all code in the supplementary material uses `Julia`’s parallel computing abilities, it will differ slightly from the examples given in the paper. For all figures, each point is the average of at least 500 replicates. We conducted the simulations in parallel on 50 Intel Xeon cores at 2.00 Ghz. All random networks were generated using the implementation of the niche model of food webs (Williams & Martinez 2000), also provided in `BioEnergeticFoodWebs`.

### Effect of carrying capacity on diversity

Starting from networks generated with the niche model with 20 species and connectance of 0.15 ± 0.01, we investigate the effect of increasing the carrying capacity of the resource (on a log scale from 0.1 to 10). We use three values of the *α*_*ij*_ parameter, ranging from 0.92 (the inter-specific competition is smaller than the intraspecific competition, which should favor coexistence), neutrally stable (intra- = inter-specific competition = 1), to 1.08 (the intraspecific competition is smaller the inter-specific competition, which should favor competitive exclusion).

We run the simulations with the default parameters (given in `?model_parameters`, and in the manual). Each simulation consists of the following code:

~~~
*# We generate a random food web*
A = nichemodel(20, 0.15)
~~~

~~~
*# This loop will keep on trying food webs
# until one with a connectance close enough
# to 0.15 is found*
while abs(BioEnergeticFoodWebs.connectance(A)-0.15)>0.01
    A = nichemodel(20, 0.15)
end
~~~

~~~
*# Prepare the simulation parameters*
for α in linspace(0.92, 1.08, 3)
  for K in logspace(-1, 1, 9)
    p = model_parameters(A, a=a,
        K=K,
        productivity=:competitive)
    *# We start each simulation with*
    *# random biomasses in ]0;1[*
        bm = rand(size(A, 1))
        *# And finally, we simulate.*
        out = simulate(p, bm, start=0,
              stop=2000, use=:ode45)
        *# And measure the output*
        diversity = foodweb_evenness(out,
                        last=1000,
                        threshold=eps())
  end
end
~~~

The results are presented in Figure 1.

### Effect of consumer-resource body-mass ratio on stability

In Figure 2, we illustrate how the effect of body-mass ratio differs between food webs with invertebrates and ectotherm vertebrate consumers.

The body-mass ratio is controlled by the parameter *Z* (field **Z** in the code), and can be changed in the following way:

~~~
scaling = logspace(-2, 4, 19) *#creates an array with 19 body-mass ratio values*
*# Prepare the simulation parameters*
p = model_parameters(A, Z=scaling[i]) *#where i is a number from 1 to 19*
~~~

Which species is an ectotherm vertebrate is controlled by the parameter `vertebrate` of `model_parameters`, which is an array of boolean (true/false) values. In order to have all consumers be ectotherm vertebrates, we use

~~~
vert = round(Bool,trophic_rank(A).>1.0)
~~~

so that for each network, we prepare the simulations with

~~~
*# Prepare the simulation parameters*
p = model_parameters(A,
      Z=scaling[i],
      vertebrates=vert)
*# where i is a number from 1 to 19, as there are 19 body-mass*
*# scaling array*
~~~

**Figure 2.**
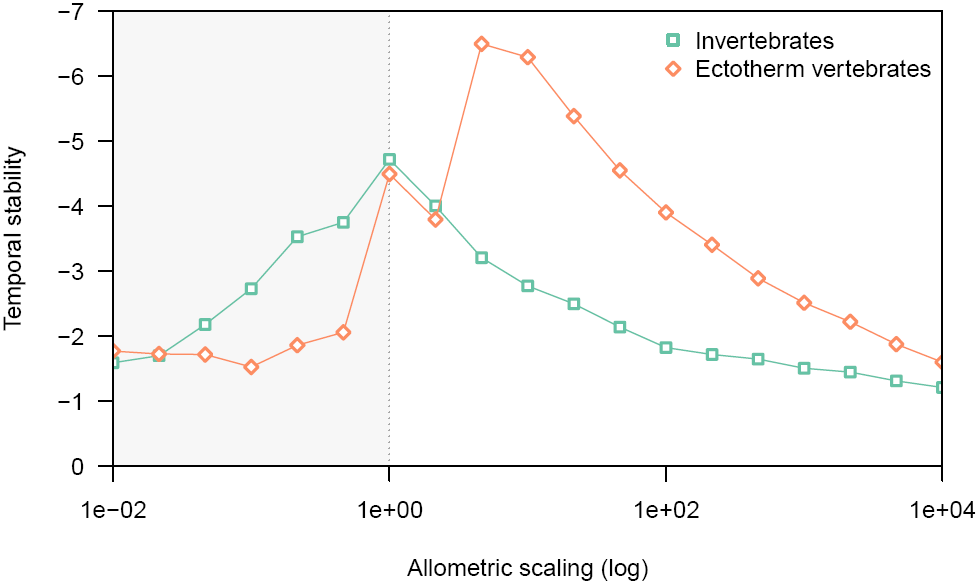
The peak of stability, in terms of allometric scaling, differs between vetebrates and invertebrates. Note that the *y* axis is reversed, since more negative values indicate less variation, and therefore more temporal stability. The shaded area represents negative scaling, *i.e.* predators are smaller than their preys.

### Effect of connectance on coexistence

We investigate the effect of connectance on species coexistence under different scenarios of inter-specific competition rates between producers (Figure 3). These simulations therefore measure how the persistence of the entire food web is affected by competition at the most basal trophic level. The persistence is used here as the measure of coexistence.

~~~
for co in vec([0.05 0.15 0.25])
  *# We generate a random food web*
   A = nichemodel(20, co)
   while abs(BioEnergeticFoodWebs.connectance(A)-co)>0.01
       A = nichemodel(20, co)
   end
   *# prepare the simulation parameters*
   for α in linspace(0.8, 1.2, 7)
     p = model_parameters(A, a=a,
     productivity=:competitive)
     bm = rand(size(A, 1))
   *# And finally, we simulate.*
     out = simulate(p, bm, start=0,
           stop=2000, use=:ode45)
     *# And measure the output*
     persistence = species_persistence(out,
                   last=1000,
                   threshold=eps())
  end
end
~~~

**Figure 3.**
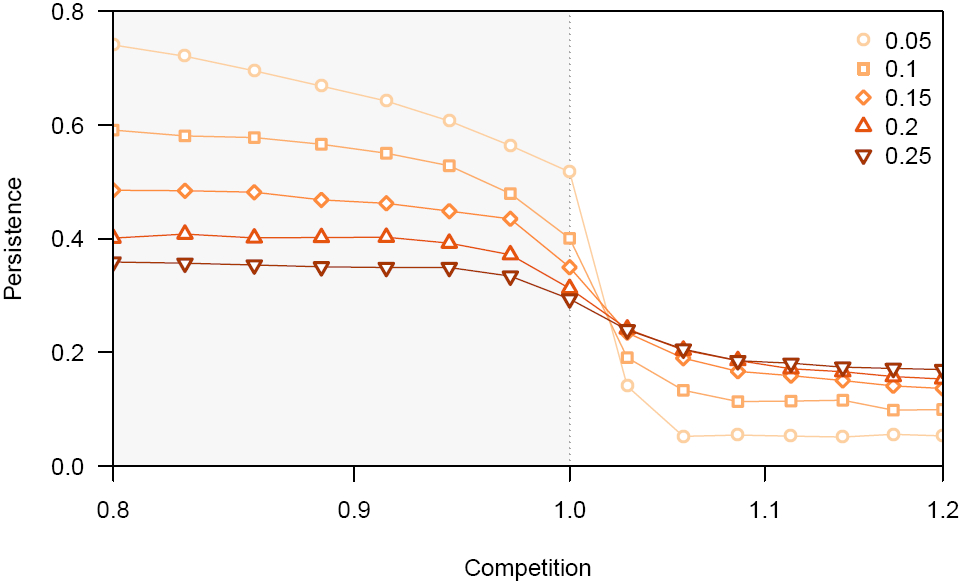
Although maximal species persistence is reached for values of inter-specific competition lower than unity, the increased trophic control at higher connectances allow coexistence even under stronger competition. The shaded area represents values of *α* smaller than unity, *i.e.* coexistence is favored.

Values of *α* larger than 0 should result in competitive exclusion in the absence of trophic interactions (Williams 2008). Indeed, this is the case when *C*_*o*_ *=* 0.05 (only a single consumer remains). Increasing connectance results in more species persisting.

## CONCLUSION

We have presented `BioEnergeticFoodWebs`, a reference implementation of the bio-energetic model applied to food webs. We provided examples that can serve as templates to perform novel simulation studies or use this model as an effective teaching tool. Because the output can be exported in a language-neutral format (JSON), the results obtained with this model can be analayzed in other languages that are currently popular with ecologists, such as `R, python`, or `MatLab`. Because we provide a general implementation that covers some of the modications made to this model over the years, there is a decreased need for individual scientists to start their own implementation, which is a both a time consuming and potentially risky endeavor.

## Acknowledgements

TP acknowledges financial support from NSERC, and an equipment grant from FRQNT. We thank the developers and maintainers of `ODE.jl`.

